# SARS CoV-2 nucleoprotein enhances the infectivity of lentiviral spike particles

**DOI:** 10.1101/2021.02.11.430757

**Authors:** Tarun Mishra, M Sreepadmanabh, Pavitra Ramdas, Amit K Sahu, Atul Kumar, Ajit Chande

## Abstract

The establishment of SARS CoV-2 spike-pseudotyped lentiviral (LV) systems has enabled the rapid identification of entry inhibitors and neutralizing agents, alongside allowing for the study of this emerging pathogen in BSL-2 level facilities. While such frameworks recapitulate the cellular entry process in ACE2+ cells, they are largely unable to factor in supplemental contributions by other SARS CoV-2 genes. To address this, we performed an unbiased ORF screen and identified the nucleoprotein (N) as a potent enhancer of spike-pseudotyped LV particle infectivity. We further demonstrate that this augmentation by N renders LV spike particles less vulnerable to the neutralizing effects of a human IgG-Fc fused ACE2 microbody. Biochemical analysis revealed that the spike protein is better enriched in virions when the particles are produced in the presence of SARS CoV-2 nucleoprotein. Importantly, this improvement in infectivity is achieved without a concomitant increase in sensitivity towards RBD binding-based neutralization. Our results hold important implications for the design and interpretation of similar LV pseudotyping-based studies.

## Introduction

The ongoing coronavirus disease 2019 (COVID-19) pandemic has provided a strong impetus for studies aimed at discovering and characterizing neutralizing antibodies or small molecule inhibitors targeted against the severe acute respiratory syndrome coronavirus 2 (SARS CoV-2). A major focus of these efforts has been the spike (S) glycoprotein of the virus, which mediates the viral entry into target cells by recognition and binding to the cell-surface receptor angiotensin-converting enzyme 2 (ACE2) ^1,2^.

Numerous studies have demonstrated that the spike protein may be employed to generate stably pseudotyped lentiviral, retroviral, and vesicular stomatitis virus particles ^3–7^. Given that live SARS CoV-2 is a biosafety level-3 agent, these pseudotyping approaches have greatly facilitated investigations undertaken within lower containment facilities. Importantly, this has also allowed for the screening and characterization of neutralizing antibodies against SARS CoV-2, as these primarily show reactivity against the spike protein ^8^. Furthermore, such pseudoviruses are deployable as platforms for the large-scale screening of small molecule inhibitors and pharmacological agents which possess therapeutic potential against COVID-19 ^9^.

In this regard, capitalizing on the interaction of the S protein with cellular ACE2, strategies have been reported which utilize chimeric ACE2 fused to the Fc region of human IgG as a potent neutralizing agent against spike-pseudotyped viruses ^10^. This arsenal now includes variations such as mutations to the catalytic region of ACE2 (thereby preventing undesirable side-effects during in vivo administration), and modified Fc domains ^10,11^. However, a common feature of these studies is that they invariably adopt pseudoviruses which are enveloped by spike, but do not incorporate contributions from any other SARS CoV-2 genes.

This point is particularly crucial in light of the manifold roles adopted by various genes of the SARS CoV-2, as has been highlighted in recent interactome studies ^12,13^. Indeed, the specific effects of these genes continue to be investigated and represent a wide diversity of specialized functions. Building on this, we hypothesized that specific genes of the SARS CoV-2 could play a key role in enhancing the infectivity of viral particles. To probe this, we undertook an unbiased screen of twenty-four ORFs, NSPs, and structural protein genes of the SARS CoV-2 using spike-pseudotyped lentiviral particles. Our observations implicated the N protein as an enhancer of viral infectivity. We further show that this enhancement of infectivity renders the viral particles less sensitive to ACE2-Immunoglobulin chimera-mediated neutralization.

## Results

A custom-designed transfer vector (**Fig. 1A**), termed pScalps Luciferase-Zsgreen, as well an ACE2-expressing HEK293T cell line (**Fig. 1B**) were generated for this study. To investigate the effect of various SARS CoV-2 genes on viral infectivity, we undertook an unbiased screen, as outlined by the schematic in **Fig. 1C**. Briefly, HEK293T producer cells were transfected with plasmids encoding the codon-optimized S protein (with a nineteen amino-acid deletion at the C terminal), the packaging plasmid psPAX2, and the pScalps Luciferase-Zsgreen reporter plasmid. Alongside this, each one of the twenty-four SARS CoV-2 genes was separately co-transfected with the above.

**Figure 1.**
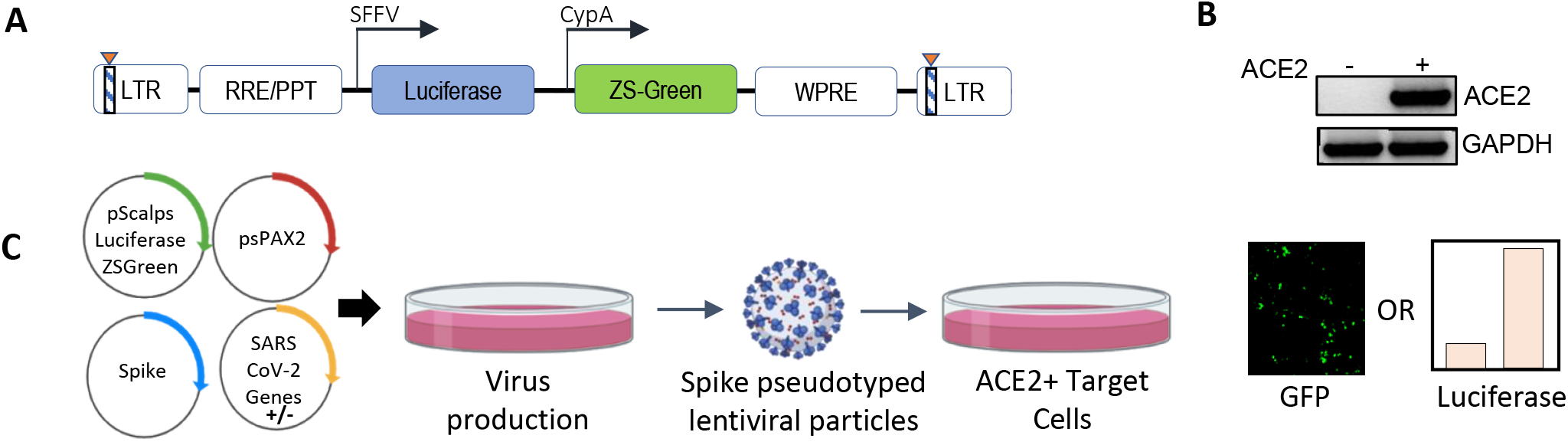
Experimental setup and constructs. (A) Schematic representation of the pScalps Luciferase Zsgreen lentiviral reporter construct. (B) ACE2 expression from the hygromycin selected HEK293T cell line transduced with a ACE2 gene carrying lentiviral vector. (C) Schematic depiction of the SARS CoV-2 genes’ screen. Transfection with an empty vector served as control. Readout from the screen was obtained in the form of firefly luciferase activity or GFP-positive cell count.

ACE2-expressing HEK293T cells were transduced with viruses produced in the presence of indicated SARS CoV-2 genes, and the extent of infectivity was gauged as a measure of both GFP-positive cell count and firefly luciferase unit’s assessment 48 hours post-infection (**Fig. 2A**). Expression of all the SARS CoV-2 genes was verified at the RNA level and the effect of this expression was also checked on the lentiviral capsid levels by western blotting from the producer cell lysates (**Fig. 2B and 2C**). Based on this screen, we identified the nucleocapsid (N) protein as an enhancer of spike-pseudovirus infectivity. We scored the effects on infectivity in conditions where the co-expression of SARS CoV-2 genes increased the infectivity without an apparent effect on the lentiviral capsid levels (**Fig. 2C**). The particle infectivity enhancement by N was also discernible with a GFP reporter suggesting this assay reliably measures transduction events (**Fig. 2D**). The infectivity-enhancement effect of N was not observed in the case of VSV-G pseudotyped lentiviral particles (**Fig. 2E**). Cumulatively, these results appear to implicate the viral N protein as an intrinsic enhancer of lentiviral spike-pseudotyped particle infectivity.

**Figure 2.**
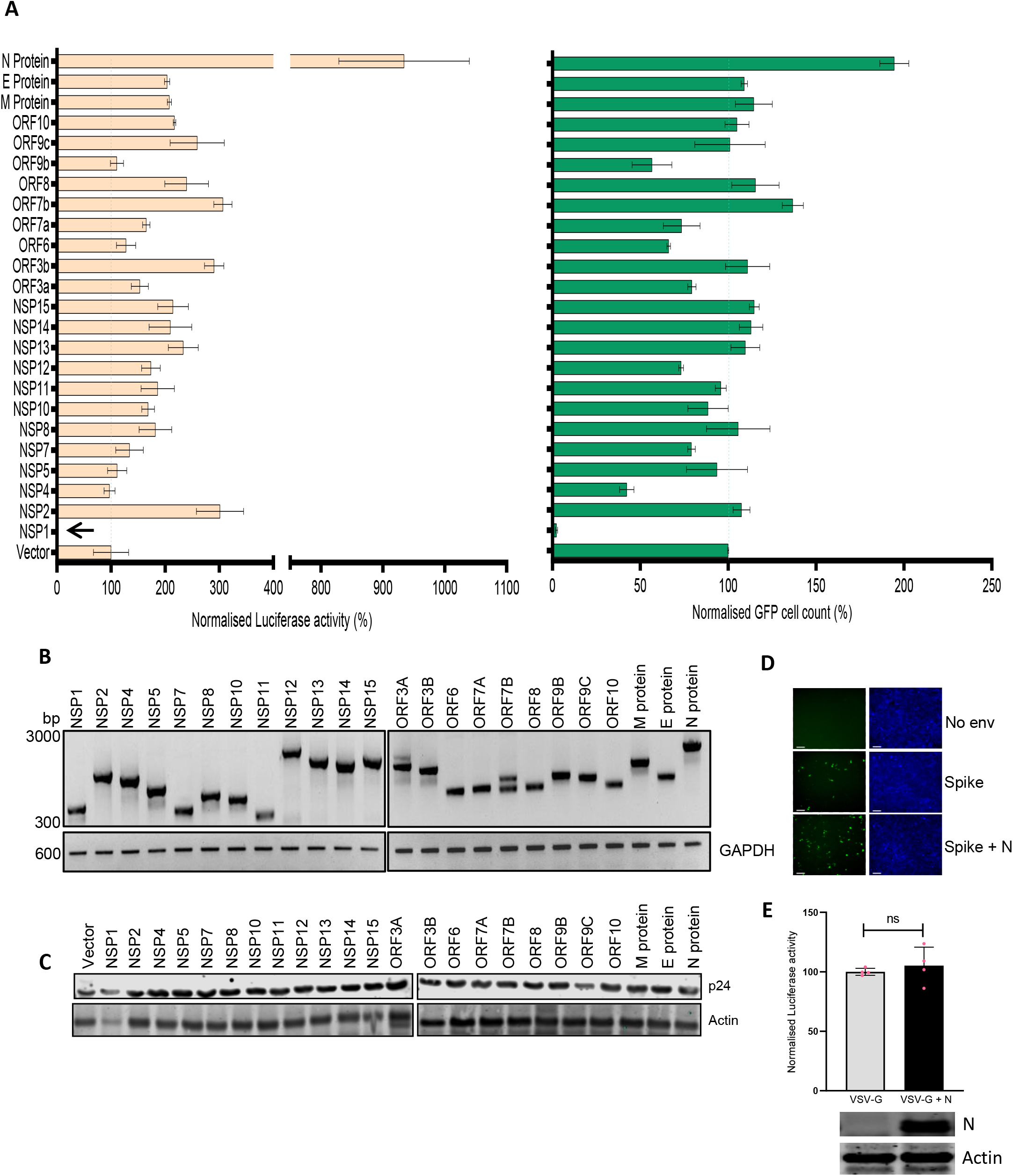
SARS CoV-2 N enhances lentiviral spike-particle infectivity. (A). Results from the screen of SARS CoV-2 genes, either by quantifying luciferase activity (left panel) or GFP positive cell count (right panel). Vector normalized values for both readout methods has been shown, such as to indicate the percentage change relative to the baseline (indicated by the dotted line) n=4 ±SD. (B) Expression of each of the SARS CoV-2 genes was verified by RT-PCR and (C) Western blotting depicting the p24 levels from the virus-producing cells. Actin served as loading control. (D) Microscopy images for the infected cells with indicated conditions (scale 100 μM). (E) Empty vector normalized Luciferase readout for VSV-G pseudotyped lentiviral particles. n=4 ±SD.

Next, we asked if such an enhancement of particle infectivity would impact the ability of neutralizing agents to block these particles from infecting host cells. Capitalizing on the interaction of the S protein with cellular ACE2, we generated a SARS CoV-2 entry receptor-based synthetic microbody consisting of a soluble ACE2 domain fused with the Fc region of human IgG (**Fig. 3A**), which also carried a C-terminal 6XHistidine tag. Extracellular expression, as well as the ability of ACE2-IgFc to dimerize under the experimental conditions that retained disulphide linkage, were detected by Western blotting (**Fig. 3B**). These experiments also confirmed the presence of an intact Histidine tag. Furthermore, we tested the specific interaction of the ACE2-IgFc with spike-pseudotyped LV particles by a pull-down experiment wherein the on-bead capture of viral particles using protein G-bound ACE2-IgFc resulted in a ~10,000-fold enrichment of spike pseudotyped viruses over VSV-G pseudotyped or non-enveloped (bald) lentiviral particles (**Fig. 3C**). This also suggests that Fc configuration remained intact after fusion with ACE2 thereby facilitating the immobilization of the microbody on the protein G magnetic beads for virion capture. In sum, these results established that the ACE2-IgFc molecule is stably expressed and demonstrates a high affinity specifically towards the S glycoprotein-laden particles.

**Figure 3.**
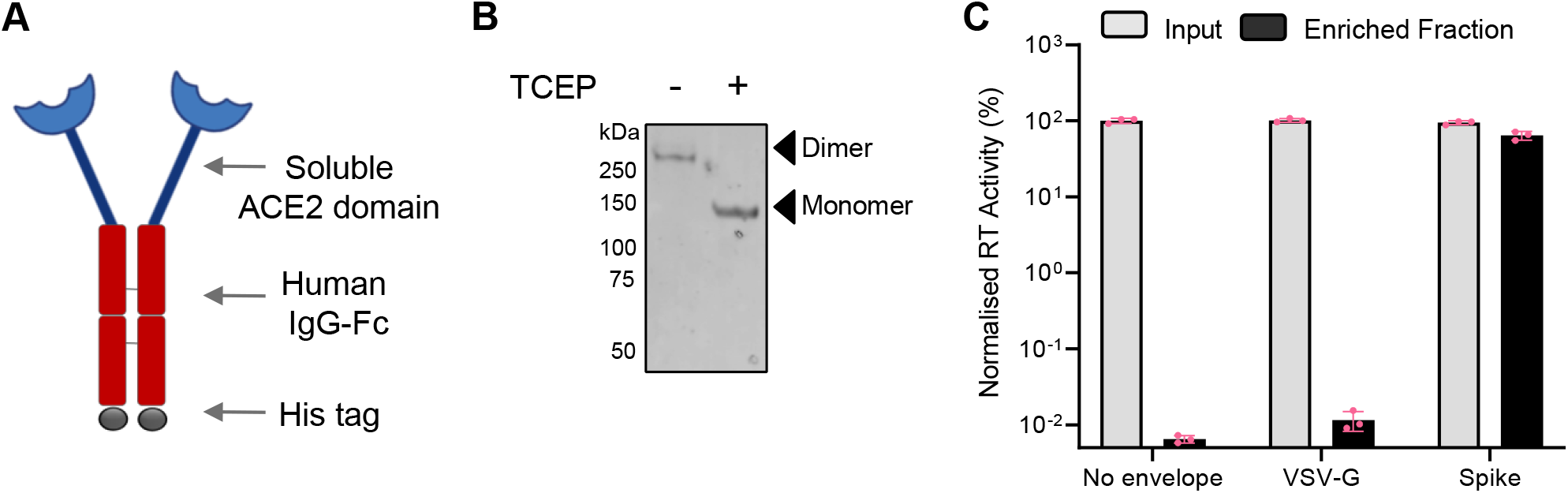
Characterization of the ACE2-IgFc microbody. (A) Schematic of the ACE2-IgFc neutralizing microbody. (B) Effect of a reducing agent (TCEP) on the migration of ACE2-IgFc on the gel. Anti-His antibody indicating monomeric vs dimeric states of the protein visualized by western blotting from the cell culture supernatant. (C) Protein G-bound ACE2-IgFC and the enrichment of virus particles. The RT activity of the input VSV-G or Spike particles was normalized to the no-envelope particles, set to 100%. n=3 ±SD.

Following this, we wished to ascertain the relative titer of ACE2-IgFc required to effectively restrict the infection of ACE2+ target cells, under the influence of the N protein. Briefly, lentiviral particles either pseudotyped with S protein (with and without co-transfection with N) or bald particles lacking envelope were generated, normalized to the RT units, incubated with ACE2-IgFc according to the indicated concentrations, and used to transduce ACE2+ HEK293T target cells. Luciferase activity assay, as a quantitative and sensitive indicator of transduction events, revealed the ability of ACE2-IgFc to impair the infectivity of spike pseudotyped viruses in a dose-dependent manner. Furthermore, results obtained herein demonstrate the requirement for at least one log higher neutralizing titer of ACE2-IgFc microbodies in the case of N-enhanced particles as opposed to those generated solely by the spike glycoprotein in order to elicit similar levels of inhibition (**Fig. 4A**). The higher requirement of neutralizing agent was not a result of higher particle counts as the inoculum was normalized to the reverse transcriptase units prior to the target cell challenge.

**Figure 4.**
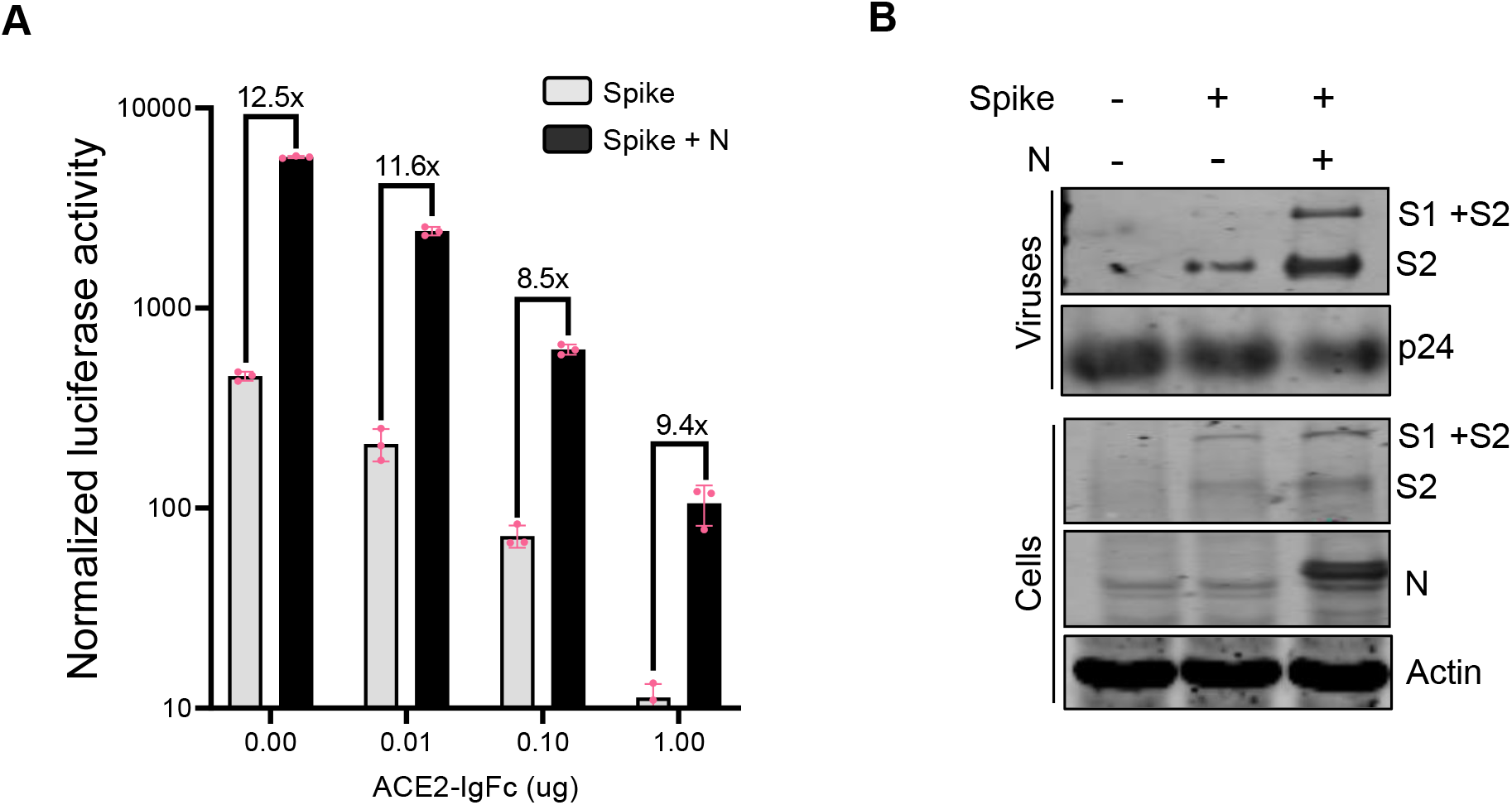
Neutralizing effects of ACE2-IgFC on the N-enhanced spike lentiviral particles and biochemical characterization of the virions (A) Neutralization assay using ACE2-IgFc pre-treated lentiviral particles with indicated amount of ACE2-IgFC (on the x-axis) for the lentiviral particles produced in the presence of indicated genes. The particles were normalized to the reverse transcriptase units obtained from the SGPERT assay before addition to target cells. Luciferase activity obtained from non-enveloped particle transduction was considered as baseline. The lines represent extent of residual infectivity (expressed as x-fold), post-treatment of ACE2-IgFc, with respect to N-exclusive particles n=3 ±SD. (B) Biochemical analysis of the virus particles and the corresponding cell lysates. The viral proteins were visualized by western blotting using antibodies raised against S2 domain of SARS CoV2 spike. The N protein and p24 were detected using Anti-Strep tag and antip24 antibody, respectively. Actin served as loading control.

Based on this, we envisaged a modification of virus particles by N that plausibly impacted the virion quality, rather than the quantity. To better understand this, we asked if N protein directly impacted the spike protein itself in order to effect infectivity enhancement. Accordingly, the viral particles produced under indicated conditions were pelleted on a sucrose cushion as described earlier ^14^. Biochemical analysis of the virus pellet and the corresponding cell lysates by Western blotting revealed that while N modestly improved the steady-state levels of spike in the producer cell lysates, the spike signal was noticeably more prominent from the virions (**Fig. 4B**). The augmentary effects of N appear to be specific in nature, given that exclusively lentiviral components do not show any apparent changes in expression. Altogether, these experiments indicated that N protein likely enhances the incorporation of spike protein into the virions, thereby improving the particle quality and its infectivity.

## Discussion

Considering the technical difficulties associated with studying live, replication-competent native SARS CoV-2, the adoption of lentiviral or retroviral pseudotyping-based systems has enabled a more widespread and accessible investigation of this pathogen. Significant advances have been achieved in recent times, ranging from optimization of pseudotyping protocols to the application of these for screening entry inhibitors ^9^. The roles of structural proteins such as M, E, and N in assisting viral assembly and particle formation have also been explored ^6^. Along similar lines, leading studies have deciphered the manifold cellular associations and interactions of various SARS CoV-2 genes, in processes as diverse as host translation inhibition, disruption of splicing, immune evasion, and protein trafficking, among others ^15–18^.

In the present study, we have evaluated the potential of twenty-four SARS CoV-2 genes to elicit superior infectivity levels as compared to what may be achieved by mere pseudotyping with the spike glycoprotein. Chief among these has been the nucleocapsid (N) protein. While a previous study has established the crucial role played by N in viral genome processing and nucleocapsid assembly ^19^, to the best of our knowledge there have been no prior reports specifically highlighting its ability to elevate lentiviral spike pseudovirus infectivity.

An interesting parallel here is the recent report that the emerging dominant D614G spike mutant is, regardless of the increased infectivity, rendered more susceptible to neutralization ^20^. In direct contrast, here we observe an alternate scenario at play - albeit within the restricted context of lentiviral pseudotypes. Our assessment of spike pseudotyped viruses generated from both N-exclusive and N-inclusive backgrounds indicates that the latter requires a higher neutralizing titer of chimeric ACE2-IgFc molecules to effectively restrict infection of ACE2+ cells. This directly implicates N as a potent enhancer of virion quality and infectivity, without an accompanying increment in vulnerability towards RBD binding interactions-based neutralizing agents. The evidence pointing towards the enrichment of spike glycoprotein within viral fractions in an N-dependent manner lends support to this hypothesis. Set against the backdrop of recent reports which highlight how specific mutations in the spike glycoprotein significantly bolster infectivity ^21^, the identification of N in this particular role may be a telling example of how the SARS CoV-2 adopts a host of varied and as-yet incompletely explored strategies to drive its pathogenesis.

In sum, our results underscore the importance of factoring in additional contributions from the other viral genes while undertaking lentiviral vector pseudotyping with the spike glycoprotein of the SARS CoV-2. While existing studies have managed to develop and validate platforms for the screening of neutralizing antibodies and inhibitory chemical compounds, we believe that our results highlight the necessity of incorporating additional genetic elements which have been shown to boost viral infectivity. Furthermore, putative therapeutic candidates currently being scrutinized in various clinical studies would benefit from lentiviral pseudotyping assays that incorporate additional virion components during the large-scale screening stage.

## Methods

### Cell culture

HEK293T (from ATCC) were cultured in DMEM containing 10% Fetal Bovine Serum (US origin certified serum, 1% penicillin-streptomycin and 20mM L-glutamine (complete medium), all obtained from Gibco. The ACE2 expressing stable HEK293T cell line was generated by transduction with lentiviral particles followed by selection with hygromycin until the entire non-transduced population was eliminated. Expression of ACE2 was verified as described below.

### Plasmids

The list of plasmids that were used in this study is provided below (refer to “Table S1: List of Plasmids”) and are available upon a reasonable request. pScalps-Luciferase-Zsgreen was generated by cloning a firefly luciferase gene that was PCR amplified with primers incorporating the XhoI/EcoRI restriction sites. The resultant PCR product was digested and ligated in the pScalps ZSgreen plasmid using identical sites and the inserts were confirmed by Sanger sequencing.

ACE2 expressing lentiviral plasmid was generated by amplifying ACE2 encoding gene from the Addgene plasmid #154987 and cloned into a modified pScalps lentiviral vector carrying the hygromycin selection marker. SARS CoV2 encoding genes (detailed below in “Table S1: List of Plasmids”) were generously provided by the Nevan Krogan Lab in the lentiviral backbone pLV-TetONE, which were further subcloned into a non-viral pcDNA based custom-designed vector for expression of these gene in transient transfection assays. After cloning, each gene of interest preserved the Strep tag in-frame at the C-terminus.

All the oligos for generation/sequencing of the plasmids used in this study were custom synthesized by Sigma Aldrich.

### Lentivirus production and quantification

In general, lentiviral particles were produced by calcium phosphate transfection of HEK293T with 8 μg of transfer vector, 6 μg of packaging plasmid (psPAX2) and 2 μg of envelope plasmid (spike-expressing plasmid or pMD2.g for virus pull-down assay). The culture medium was replaced at 16h-post transfection. Lentiviral vector-containing supernatant was collected 48h after transfection and was centrifuged and filtered through 0.22 μM syringe filters. The infectivity assay was performed after normalizing reverse transcriptase (RT) units obtained from an SGPERT assay as described earlier ^14^. Briefly, the target cells were infected in quadruplicates (or at least triplicate) with up to 125-fold dilutions and the infectivity was acquired from the dilutions in the linear range as reported previously ^14^. Expression of GFP or Luciferase as a quantitative measure of infection was acquired using Spectramax plate reader or CX7 High-content imaging platform (CellInsight CX7 High Content Screening platform, ThermoFisher Scientific and SpectramaxI3X, Molecular Devices, USA).

The SARS CoV-2 genes’ screen was performed by producing spike pseudotyped lentiviruses from HEK293T cells that were seeded in 12-well plate. Next day cells were transfected with pScalps luciferase ZSgreen (800 ng), psPax2 (600 ng), spike expression plasmid (200 ng), and either vector control or ORF carrying vector (300 ng) by the calcium phosphate method. Culture medium was replaced with fresh medium 16h after transfection. Spike pseudotyped viruses were collected after 48h of transfection. Supernatant was collected and centrifuged at 1200Xg for 5 min and filtered using a 0.22 μM syringe filter.

Spike pseudotyped viruses were then used to transduce ACE2 expressing HEK293T cells to check the infectivity level by either by GFP positive cell count and or by Luciferase assay.

To quantify the RT units, 5 μl viruses were lysed in a 5 μl lysis buffer and diluted with a core buffer to make volume 100 μl. A 10 μl diluted viral lysate was mixed with an equal volume of 2X reaction buffer and the SGPERT assay was performed.

For immunoblotting of proteins incorporated into lentiviral particles, the lentiviral particles-containing supernatant was concentrated on sucrose cushion by ultracentrifugation at 50,000Xg for 2 hours at 4◻ ^14^. Viral pellet obtained was resuspended in the Laemmli buffer containing 10mM TCEP as a reducing agent.

### SARS CoV-2 Genes and ACE2 expression

To check the expression of ACE2 in ACE2-transduced HEK293T cells we isolated the RNA from HEK293T and ACE2+ HEK293T cells using Trizol and synthesized cDNA from 1ug of RNA using oligoDT primers obtained from both the cells. To quantify the level of ACE2 expression, qPCR was performed using ACE2 specific primers 5’-GGGATCAGAGATCGGAAGAA-3’ forward and 5’-AGGAGGTCTGAACATCATC-3’ reverse and GAPDH as control with primers 5’-TGGAGAAGGCTGGGGCTCATTTGCA-3’ forward and 5’-CATACCAGGAAATGAGCTTGACAA-3’.

RT-PCR for SARS CoV-2 genes from the producer HEK293T cells was performed using the forward primer 5’-TCCTACCCTCGTAAAGAATTC-3’ and reverse primer 5’-TCCGGACTTTTCAAACTGCGGATGT-3’ with the respective cDNA as template.

### Luciferase assay

Luciferase assay was performed in 96-well plates to quantify level of transduction by spike pseudotyped viral particles. Equal numbers of ACE2+ HEK293T cells were seeded in 96-well plates (10,000 cells/well) 24h before transduction and incubated with the various pseudotyped viruses for 48h. GFP cell count was scored using the SpectraMax i3X system (Molecular Devices). Media was aspirated from each well after 48h and Cells were lysed by treatment with 100 μl lysis buffer for 20 minutes at room temperature. Luciferase readings were measured using SpectraMax i3X by injecting 50 μl of substrate buffer in 50 μl of cell lysate in 96-well white plates. Specifics of buffer composition may be found below in the section titled “Buffers and common reaction mixtures”.

### Western blotting

For Western blot-based analysis, samples were prepared in a 4X Laemmli buffer, boiled for 5 min at 95°C, and run on either 8% or 12.5% tricine gels for electrophoresis depending upon the molecular weight range being detected by this method. Following this, gels were electro blotted on the PVDF membrane (Immobilon-FL, Merck-Millipore). Blocking of membranes was carried out by incubation with either a 5% BSA solution or the proprietary Odyssey Blocking Buffer (LI-COR Biosciences) for one hour, followed by both primary and secondary antibody incubations for one hour each at room temperature, each of which were followed by three washes for five minutes.

Detection of p24, beta actin, SARS CoV-2 spike glycoprotein, SARS CoV-2 genes, and ACE2-IgFc was carried out using mouse anti-p24 (NIH ARP), rabbit anti-beta actin (LI-COR Biosciences, Cat# 926-42210, RRID:AB_1850027), mouse anti-spike (Cat#ZMS1076, Sigma Aldrich), mouse anti-Strep (Qiagen, Cat# 34850), and mouse anti-Histidine (Invitrogen, Cat# MA1-21315), respectively, as primary antibodies. Secondary antibodies used were either IR dye 680 goat anti-mouse, IR dye 800 goat mouse, or IR dye 800 goat anti-rabbit (LI-COR Biosciences Cat# 925-68070, RRID:AB_2651128, and LI-COR Biosciences Cat# 925-32211, RRID:AB_2651127).

### Generation of a plasmid expressing synthetic ACE2-IgFc microbody

Genomic DNA isolated from HEK293T cells using the Macherey Nagel Nucleospin kit (#740952.50) was used as a template for PCR amplifying the Fc region of human IgG. The forward primer 5’-CAGCACCTGAACTCCTGGGGGGACCG-3’ and reverse primer 5’-CCTTTGGCTTTGGAGATGGTTTTC-3’ was used to amplify the first exon encoding the Fc region. The forward primer 5’-AGGGCAGCCCCGAGAACCACAGGTG-3’ and reverse primer 5’-TTTACCCGGAGACAGGGAGAGGCT-3’ was used to amplify the second exon encoding the Fc region. Both fragments were individually amplified further using the combination of forward and reverse primers as 5’-GAAAACCATCTCCAAAGCCAAAGGGCAGCCCCGAGAACCACAGGTG-’3’ and 5’-TATATATTCTAGATTAATGGTGATGGTGATGATGGCCGCCTTTACCCGGAGACAGGGA-3’ for the first exon’s amplicon and 5’-ATATATCTCGAGGACAAAACTCACACATGCCCACCGTGCCCAGCACCTGAACTCCTG-3’ and 5’-CCTTTGGCTTTGGAGATGGTTTTC-3’ for the second exon’s amplicon, respectively.

Finally, both these fragments were combined using the forward primer 5’-ATATATCTCGAGGACAAAACTCACACATGCCCACCGTGCCCAGCACCTGAACTCCTG-3’ and reverse primer 5’-TATATATTCTAGATTAATGGTGATGGTGATGATGGCCGCCTTTACCCGGAGACAGGGA-3’ to generate the final Fc fragment, which was cloned into pTZ57R.

Addgene (#154987) was used as template for ACE2 amplification, using the forward primer 5’-GAACAAGAATTCTTTTGTGGGA-3’ and reverse primer 5’-TTTGTCCTCGAGGGAAACAGGGGGCTGGTTAG-3’ a 421 bp fragment of the same was amplified and cloned into pTZ57R.

Using an EcoRI/XhoI digestion, the ACE2 fragment was combined upstream of the Fc, and the combined construct was ligated into the ACE2 containing plasmid using a EcoRI/XbaI digestion to yield the final ACE2-IgFc construct. The construct was verified with Sanger sequencing (refer to “Table S2: Sequences of ACE2-IgFc”).

### Biochemical characterization of ACE2-IgFc microbody

HEK293T cells were transfected with the plasmid encoding ACE2-IgFc (5 μg plasmid in a 35 mm dish). As a control, 5 μg of pcDNA3.1BS(-) was also transfected. Media was changed twelve hours post-transfection and fresh DMEM (without FBS) was added to the dishes. Culture media was collected both 48- and 72-hours post-transfection, and samples were prepared in either 8% or 2% SDS-containing 4X Laemmli buffer (with and without TCEP added, respectively) for SDS-PAGE. 8% tricine gels were run, and analyzed using Western Blotting. The amount of ACE2 present in the supernatant was determined by comparison with standard concentrations of pure BSA as ascertained by band density analysis following silver staining of SDS-PAGE gels. For Western Blotting, the proteins were transferred to a PVDF membrane, blocked in a 5% BSA solution in TBS, and primary mouse-derived anti-Histidine antibodies were incubated with the blot in a 1:4000 dilution, followed by goat-derived anti-mouse antibodies in a 1:5000 dilution.

### Pull-down assay using ACE2-IgFc

A 30 μl of protein-G Dynabeads (Thermofisher Scientific Cat #10003D) were first equilibrated by washing with 10% FBS-containing DMEM. Following this, these were incubated with gentle mixing by inversion for 15 minutes with 100 μl of ACE2-IgFc containing supernatant (equivalent of 10 μg/ml). This was split into three equal fractions of 10 μl each. The beads-containing mixtures then were immobilized on a magnetic rack, supernatant was aspirated off, and replaced with 100 μl (10mU equivalent/mL RT) of culture supernatant carrying either bald viruses or viruses pseudotyped with VSV-G or Spike glycoproteins. Mixing was carried out for a period of 15 minutes, following which these were placed back on the magnetic rack, supernatant was aspirated, and the beads washed twice with SGPERT Core Buffer - first with 500 μl and then with 100 μl. Finally, 20 μl of RT Lysis Buffer was added and the reaction was incubated at room temperature before mixing with TritonX diluted 10-fold using core buffer. The resulting supernatant was collected after immobilizing the beads and processed for an SGPERT Assay.

### Neutralization Assay

To quantify the amount of ACE2-IgFc required to inhibit the infection in ACE2+ HEK293T cells, we treated 100 μl of spike pseudotyped virus (normalised to RT units) with a variable amount of ACE2-IgFc containing medium. The mixture was incubated for fifteen minutes at room temperature, following which each sample was used to transduce ACE2+ HEK293T cells seeded in 96-well plates. Image acquisition (CellInsight CX7 High Content Screening platform, ThermoFisher Scientific) GFP cell count (SpectramaxI3 Molecular Devices), and luciferase assay were performed 48 hours post-transduction, in the enlisted order.

### Software

All graphs were generated using GraphPad Prism (version 9.0). Specific portions of images were produced using Biorender.

## Acknowledgment

This work was supported by the intramural funds, a Department of Biotechnology (DBT) grant (BT/PR26013/GET/119/191/2017), and the Wellcome Trust/DBT India Alliance Fellowship [grant number IA/I/18/2/504006 awarded to AC]. TM and PR are supported by a fellowship from the MHRD. AC is a recipient of the Innovative Young Biotechnologist Award from the DBT. Authors are grateful to Nevan Krogan, Massimo Pizzato, Jeremy Luban, Sonja Best, Raffaele De Francesco, Didier Trono, Sunando Datta, and the NIH AIDS Reagent Program for the reagents and cell lines.

