## Supplemental table 1 & 2 for "SARS CoV-2 nucleoprotein enhances the infectivity of lentiviral spike particles"

*for*

**Table S1: List of Plasmids**

| S. No. | Plasmid Name | Purpose | Notes | Source |
|---|---|---|---|---|
| 1. | pScalps Luciferase Zsgreen | Expression of firefly luciferase and Zoanthus green fluorescent protein | Lentiviral vector with luciferase gene under SFFV promoter and Zsgreen under Cyclophilin promoter | This study |
| 2. | Spike del19 codon-optimized in pcDNA3.1(-) | Expression of mammalian codon optimized SARS CoV-2 spike glycoprotein | A nineteen amino acid deletion at the C terminal | Addgene (#155297), from Raffaele De Francesco's lab |
| 3. | psPAX2 | Lentiviral packaging plasmid | Packaging vector | Addgene (#12260) from Didier Trono's lab |
| 4. | pMD2.G | Expression of VSV-G glycoprotein | Envelope plasmid | Addgene (#12259) from Didier Trono's lab |
| 5. | pBJ5-VSV-G HA | Expression of HA-tagged VSV-G glycoprotein | Envelope plasmid | Gifted by Prof. Massimo Pizzato |
| 6. | pcDNA3.1BS(-) | Control for transfection and vector backbone for cloning | Mammalian expression vector | This study |
| 7. | pWPI-IRES-Puro-Ak-ACE2-TMPRSS2 | Source for human ACE2 | Puromycin resistance marker | Addgene (#154987) from Sonja Best's lab |

| | | | | |
|---|---|---|---|---|
| 8. | ACE2 in pScalps Hygro | Establishment of ACE2+ cell line | | This study |
| 9. | ACE2-IgFc | Expression of ACE2-IgFc construct | 6XHistidine tag at the C-terminus | This study |
| 10. | Nsp1 in pcDNA3.1BS(-) | Expression of SARS-CoV-2's non-structural protein 1 | C-terminal 2X Strep Tag | This study |
| 11. | Nsp2 in pcDNA3.1BS(-) | Expression of SARS-CoV-2's non-structural protein 2 | C-terminal 2X Strep Tag | This study |
| 12. | Nsp4 in pcDNA3.1BS(-) | Expression of SARS-CoV-2's non-structural protein 4 | C-terminal 2X Strep Tag | This study |
| 13. | Nsp5 in pcDNA3.1BS(-) | Expression of SARS-CoV-2's non-structural protein 5 | C-terminal 2X Strep Tag | This study |
| 14. | Nsp7 in pcDNA3.1BS(-) | Expression of SARS-CoV-2's non-structural protein 7 | C-terminal 2X Strep Tag | This study |
| 15. | Nsp8 in pcDNA3.1BS(-) | Expression of SARS-CoV-2's non-structural protein 8 | C-terminal 2X Strep Tag | This study |
| 16. | Nsp10 in pcDNA3.1BS(-) | Expression of SARS-CoV-2's non-structural protein 10 | C-terminal 2X Strep Tag | This study |
| 17. | Nsp11 in pcDNA3.1BS(-) | Expression of SARS-CoV-2's non-structural protein 1 | C-terminal 2X Strep Tag | This study |
| 18. | Nsp12 in pcDNA3.1BS(-) | Expression of SARS-CoV-2's non-structural protein 12 | C-terminal 2X Strep Tag | This study |
| 19. | Nsp13 in pcDNA3.1BS(-) | Expression of SARS-CoV-2's non-structural protein 13 | C-terminal 2X Strep Tag | This study |
| 20. | Nsp14 in pcDNA3.1BS(-) | Expression of SARS-CoV-2's non-structural protein 14 | C-terminal 2X Strep Tag | This study |

| | | | | |
|---|---|---|---|---|
| 21. | Nsp15 in pcDNA3.1BS(-) | Expression of SARS-CoV-2’s non-structural protein 15 | C-terminal 2X Strep Tag | This study |
| 22. | ORF3a in pcDNA3.1BS(-) | Expression of SARS-CoV-2’s open reading frame 3a | C-terminal 2X Strep Tag | This study |
| 23. | ORF3b in pcDNA3.1BS(-) | Expression of SARS-CoV-2’s open reading frame 3b | C-terminal 2X Strep Tag | This study |
| 24. | ORF6 in pcDNA3.1BS(-) | Expression of SARS-CoV-2’s open reading frame 6 | C-terminal 2X Strep Tag | This study |
| 25. | ORF7a in pcDNA3.1BS(-) | Expression of SARS-CoV-2’s open reading frame 7a | C-terminal 2X Strep Tag | This study |
| 26. | ORF7b in pcDNA3.1BS(-) | Expression of SARS-CoV-2’s open reading frame 7b | C-terminal 2X Strep Tag | This study |
| 27. | ORF8 in pcDNA3.1BS(-) | Expression of SARS-CoV-2’s open reading frame 8 | C-terminal 2X Strep Tag | This study |
| 28. | ORF9b in pcDNA3.1BS(-) | Expression of SARS-CoV-2’s open reading frame 9b | C-terminal 2X Strep Tag | This study |
| 29. | ORF9c in pcDNA3.1BS(-) | Expression of SARS-CoV-2’s open reading frame 9c | C-terminal 2X Strep Tag | This study |
| 30. | ORF10 in pcDNA3.1BS(-) | Expression of SARS-CoV-2’s open reading frame 10 | C-terminal 2X Strep Tag | This study |
| 31. | M protein in pcDNA3.1BS(-) | Expression of SARS-CoV-2’s membrane protein | C-terminal 2X Strep Tag | This study |
| 32. | E protein in pcDNA3.1BS(-) | Expression of SARS-CoV-2’s envelope protein | C-terminal 2X Strep Tag | This study |
| 33. | N protein in pcDNA3.1BS(-) | Expression of SARS-CoV-2’s nucleocapsid protein | C-terminal 2X Strep Tag | This study |

| | | | | |
|---|---|---|---|---|
| 34. | pScalps Hygro | Lentiviral vector backbone for selection of transduced cells by hygromycin | Hygromycin resistance marker and an MCS for expression under SFFV promoter | This study |
| 35. | pTZ57R | Scaffold for cloning of ACE2-IgFc | Cloning vector | Fermentas/Thermo Scientific |

**Table S2: Sequences of ACE2-IgFc:**

| | |
|---|---|
| Nucleotide sequence of ACE2-IgFc: | 5'-<br>ATGTCAAGCTCTTCCTGGCTCCTTCTCAGCCTTGTTGCTGTAACTGCTGC<br>TCAGTCCACCATTGAGGAACAGGCCAAGACATTTTTGGACAAGTTTAACC<br>ACGAAGCCGAAGACCTGTTCTATCAAAGTTCACTTGCTTCTTGGAATTATA<br>ACACCAATATTACTGAAGAGAATGTCCAAAACATGAATAATGCTGGGGAC<br>AAATGGTCTGCCTTTTTAAAGGAACAGTCCACACTTGCCCAAATGTATCC<br>ACTACAAGAAATTCAGAATCTCACAGTCAAGCTTCAGCTGCAGGCTCTTC<br>AGCAAAATGGGTCTTCAGTGCTCTCAGAAGACAAGAGCAAACGGTTGAA<br>CACAATTCTAAATACAATGAGCACCATCTACAGTACTGGAAAAGTTTGTAA<br>CCCAGATAATCCACAAGAATGCTTATTACTTGAACCAGGTTTGAATGAAAT<br>AATGGCAAACAGTTTAGACTACAATGAGAGGCTCTGGGCTTGGGAAAGC<br>TGGAGATCTGAGGTCGGCAAGCAGCTGAGGCCATTATATGAAGAGTATG<br>TGGTCTTGAAAAATGAGATGGCAAGAGCAAATCATTATGAGGACTATGGG<br>GATTATTGGAGAGGAGACTATGAAGTAAATGGGGTAGATGGCTATGACTA<br>CAGCCGCGGCCAGTTGATTGAAGATGTGGAACATACCTTTGAAGAGATTA<br>AACCATTATATGAACATCTTCATGCCTATGTGAGGGCAAAGTTGATGAAT<br>GCCTATCCTTCCTATATCAGTCCAATTGGATGCCTCCCTGCTCATTTGCTT<br>GGTGATATGTGGGGTAGATTTTGGACAAATCTGTACTCTTTGACAGTTCC<br>CTTTGGACAGAAACCAAACATAGATGTTACTGATGCAATGGTGGACCAGG<br>CCTGGGATGCACAGAGAATATTCAAGGAGGCCGAGAAGTTCTTTGTATCT<br>GTTGGTCTTCCTAATATGACTCAAGGATTCTGGGAAAATTCCATGCTAAC<br>GGACCCAGGAAATGTTCAGAAAGCAGTCTGCCATCCCACAGCTTGGGAC<br>CTGGGGAAGGGCGACTTCAGGATCCTTATGTGCACAAAGGTGACAATGG<br>ACGACTTCCTGACAGCTCATCATGAGATGGGGCATATCCAGTATGATATG<br>GCATATGCTGCACAACCTTTTCTGCTAAGAAATGGAGCTAATGAAGGATT<br>CCATGAAGCTGTTGGGGAAATCATGTCACTTTCTGCAGCCACACCTAAGC<br>ATTTAAAATCCATTGGTCTTCTGTCACCCGATTTTCAAGAAGACAATGAAA<br>CAGAAATAAACTTCCTGCTCAAACAAGCACTCACGATTGTTGGGACTCTG<br>CCATTTACTTACATGTTAGAGAAGTGGAGGTGGATGGTCTTTAAAGGGGA<br>AATTCCCAAAGACCAGTGGATGAAAAAGTGGTGGGAGATGAAGCGAGAG<br>ATAGTTGGGGTGGTGGAACCTGTGCCCCATGATGAAACATACTGTGACC |

<table>
<tr><td></td><td>CCGCATCTCTGTTCCATGTTTCTAATGATTACTCATTCATTCGATATTACA<br>CAAGGACCCTTTACCAATTCCAGTTTCAAGAAGCACTTTGTCAAGCAGCT<br>AAACATGAAGGCCCTCTGCACAAATGTGACATCTCAAACTCTACAGAAGC<br>TGGACAGAAACTGTTCAATATGCTGAGGCTTGGAAAATCAGAACCCTGGA<br>CCCTAGCATTGGAAAATGTTGTAGGAGCAAAGAACATGAATGTAAGGCCA<br>CTGCTCAACTACTTTGAGCCCTTATTTACCTGGCTGAAAGACCAGAACAA<br>GAATTCTTTTGTGGGATGGAGTACCGACTGGAGTCCATATGCAGACCAAA<br>GCATCAAAGTGAGGATAAGCCTAAAATCAGCTCTTGGAGATAAAGCATAT<br>GAATGGAACGACAATGAAATGTACCTGTTCCGATCATCTGTTGCATATGC<br>TATGAGGCAGTACTTTTTAAAAGTAAAAAATCAGATGATTCTTTTTGGGGA<br>GGAGGATGTGCGAGTGGCTAATTTGAAACCAAGAATCTCCTTTAATTTCT<br>TTGTCACTGCACCTAAAAATGTGTCTGATATCATTCCTAGAACTGAAGTTG<br>AAAAGGCCATCAGGATGTCCCGGAGCCGTATCAATGATGCTTTCCGTCT<br>GAATGACAACAGCCTAGAGTTTCTGGGGATACAGCCAACACTTGGACCT<br>CCTAACCAGCCCCCTGTTTCCCTCGAGGACAAAACTCACAAATGCCCAC<br>CGTGCCCAGCACCTGAACTCCtGGGGGGACCGTCAGTCTTCCTCTTCCC<br>CCCAAAACCCAAggAtaCCCTTATGATTTCCCGGACCCCTGAGGTCACGTG<br>CGTGGTGGTGGACGTGAGCCACGAAGACCCCGAGGTCCAGTTCAAGTG<br>GTACGTGGACGGCGTGGAGGTGCATAATGCCAAGACAAAGCTGCGGGA<br>GGAGCAGTACAACAGCACGTTCCGTGTGGTCAGCGTCCTCACCGTCCTG<br>CACCAGGACTGGCTGAACGGCAAGGAGTACAAGTGCAAGGTCTCCAACA<br>AAGCCCTCCCAGCCCCCATCGAGAAAACCATCTCCAAAGCCAAAGGGCA<br>GCCCCGAGAACCACAGGTGTACACCCTGCCCCCATCCCGGGATGAGCT<br>GACCAAGAACCAGGTCAGCCTGACCTGCCTGGTCAAAGGCTTCTATCCC<br>AGCGACATCGCCGTGGAGTGGGAGAGCAATGGGCAGCCGGAGAACAAC<br>TACAAGACCACGCCTCCCGTGCTGGACTCCGACGGCTCCTTCTTCCTCT<br>ACAGCAAGCTCACCGTGGACAAGAGCAGGTGGCAGCAGGGGAACGTCT<br>TCTCATGCTCCGTGATGCATGAGGGTCTGCACAACCACTACACGCAGAA<br>GAGCCTCTCCCTGTCTCCGGGTAAAGGCGGCCATCATCACCATCACCAT<br>TAA-3’</td></tr>
<tr><td>Translation of ACE2-IgFc:</td><td>MSSSSWLLLSLVAVTAAQSTIEEQAKTFLDKFNHEAEDLFYQSSLASWNYNT<br>NITEENVQNMNNAGDKWSAFLKEQSTLAQMYPLQEIQNLTVKLQLQALQQN<br>GSSVLSEDKSKRLNTILNTMSTIYSTGKVCNPDNPQECLLLEPGLNEIMANSL<br>DYNERLWAWESWRSEVGKQLRPLYEEYVVLKNEMARANHYEDYGDYWRG<br>DYEVNGVDGYDYSRGQLIEDVEHTFEEIKPLYEHLHAYVRAKLMNAYPSYIS<br>PIGCLPAHLLGDMWGRFWTNLYSLTVPFGQKPNIDVTDAMVDQAWDAQRIF<br>KEAEKFFVSVGLPNMTQGFWENSMLTDPGNVQKAVCHPTAWDLGKGDFRI<br>LMCTKVTMDDFLTAHHEMGHIQYDMAYAAQPFLLRNGANEGFHEAVGEIMS<br>LSAATPKHLKSIGLLSPDFQEDNETEINFLLKQALTIVGTLPFTYMLEKWRWM<br>VFKGEIPKDQWMKKWWEMKREIVGVVEPVPHDETYCDPASLFHVSNDYSFI<br>RYYTRTLYQFQFQEALCQAAKHEGPLHKCDISNSTEAGQKLFNMLRLGKSE<br>PWTLALENVVGAKNMNVRPLLNYFEPLFTWLKDQNKNSFVGWSTDWSPYA<br>DQSIKVRISLKSALGDKAYEWNDNEMYLFRSSVAYAMRQYFLKVKNQMILFG<br>EEDVRVANLKPRISFNFFVTAPKNVSDIIPRTEVEKAIRMSRSRINDAFRLNDN</td></tr>
</table>

| | |
|---|---|
| | SLEFLGIQPTLGPPNQPPVSLEDKTHKCPPCPAPELLGGPSVFLFPPKPKDTL MISRTPEVTCVVVDVSHEDPEVQFKWYVDGVEVHNAKTKLREEQYNSTFRV VSVLTVLHQDWLNGKEYKCKVSNKALPAPIEKTISKAKGQPREPQVYTLPPS RDELTKNQVSLTCLVKGFYPSDIAVEWESNGQPENNYKTTPPVLDSDGSFFL YSKLTVDKSRWQQGNVFSCSVMHEGLHNHYTQKSLSLSPGKGGHHHHHH |
| Legend: | N-terminal human ACE2 sequence; Modified C-terminal soluble ACE2 Fragment; Cloning sites; IgG Fc Fragment; 6XHistidine tag |
